# Influences of age and pubertal status on number of perineuronal nets in the rat medial prefrontal cortex

**DOI:** 10.1101/2020.01.31.929257

**Authors:** Carly M. Drzewiecki, Jari Willing, Janice M. Juraska

**Author notes:** Corresponding Author Janice M. Juraska, Department of Psychology, University of Illinois, 603 E Daniel St, Champaign, IL 61820.

## Abstract

The prefrontal cortex (PFC) is a late developing region of the cortex, and its protracted maturation during adolescence may confer a period of plasticity. Closure of critical, or sensitive, periods in sensory cortices coincides with perineuronal net (PNN) expression, leading to enhanced inhibitory function and synaptic stabilization. PNN density is known to increase across adolescence in the male rodent medial PFC (mPFC). However, the trajectory of female PNN development has not been explored nor has the potential role of pubertal onset in PNN expression. Here, we examined rats at four time points spanning adolescent development to quantify the number of PNNs in the mPFC, as well as the total volume of the prefrontal white matter. Additionally, because puberty coincides with broad behavioral and neuroanatomical changes, we collected tissue from age-matched pre- and post- pubertal siblings within a litter. Results indicate that both males and females show an increase the total number of mPFC PNNs and in white matter under the mPFC between postnatal day (P) 30 and P60. Male puberty did not affect PNNs, while female pubertal onset led to an abrupt decrease in the total number of PNNs that persisted through mid-adolescence before increasing at P60. This decrease in female rats may indicate a difference in timing of maximal plasticity between the sexes during adolescence.

## Introduction

Critical, or sensitive, periods are windows of heightened plasticity where the developing brain is especially responsive to sensory stimuli (Hensch, 2005). Neuronal circuits can be permanently shaped by experience during these sensitive windows, and sensory deprivation during this time can have long-lasting implications for brain structure and function. Critical periods for cortical development have been well-established in multiple species across several brain regions, including the visual and somatosensory cortices (Fox, 1992; Hensch, 2004; Wiesel & Hubel, 1970). In the visual cortex, they begin with the appearance of GABAergic parvalbumin (PV) interneurons, which enhance inhibitory signaling and allow for increased plasticity (Hensch, 2005). The closure of the critical period coincides with perineuronal net (PNN) development (Pizzorusso et al., 2002).

PNNs are specialized components of the extracellular matrix found throughout the central nervous system (Celio et al., 1998; Seeger et al., 1994). They are composed of an aggregation of chondroitin sulfate proteoglycans (CSPGs) called the lecticans, which include aggrecan, brevican, versican, and neurocan (Yamaguchi, 2000). The CSPGs interact with hyaluronan, link proteins and tenascin-R to form mesh-like structures surround the soma and proximal dendrites of neurons. PNNs primarily surround PV cells, where they act as a cation buffer to enhance the excitability of the fast-spiking interneurons (Balmer, 2016; Härtig et al., 1999). PNNs also contribute synaptic stability and create a barrier that prevents the formation of new synaptic contacts while stabilizing existing ones (Härtig et al., 1992). As such, the development of PNNs coincides with the closure of sensitive periods (Carulli et al., 2010). Degradation of PNNs can restore juvenile levels of ocular dominance plasticity in the adult visual cortex (Pizzorusso et al., 2002) and increase synaptic plasticity in the perirhinal cortex (Romberg et al., 2013). Given their role in modulating neuroplasticity, PNNs in both cortical and non-cortical brain regions have been implicated in a wide range of behavioral functions, including drug seeking behaviors, reversal learning, and fear conditioning (Blacktop & Sorg, 2019; Gogolla et al., 2009; Happel et al., 2014; Slaker et al., 2015).

It should be added that there are other potential contributors that influence the closure of sensitive periods such as actions of myelin derived-Nogo at the Nogo-66 receptor (NgR) that prevent axon outgrowth (Akbik et al., 2013; McGee & Strittmatter, 2003). Within the visual cortex, increasing myelination coincides with the closure of the sensitive period, and NgR null mice show enhanced plasticity in adulthood (McGee et al., 2005). The presence of this receptor has also been shown to limit plasticity in adulthood in the somatosensory cortex, amygdala, and mPFC (Bhagat et al., 2015; Jitsuki et al., 2016; Yang et al., 2012).

Though sensitive periods have been well-studied in the juveniles, more recent work suggests that adolescence remains a time of heightened neural plasticity, particularly in the late-developing prefrontal cortex (PFC) (Fuhrmann et al., 2015; Selemon, 2013). In humans, a multitude of studies have demonstrated that the frontal cortex undergoes protracted anatomical development across adolescence (Gogtay et al., 2004; Lenroot & Giedd, 2006; Sowell et al., 1999), and much of this maturation corresponds with increased executive functioning and behavioral inhibition capabilities (Anderson et al., 2001; Taylor et al., 2013). Pubertal onset is a hallmark of adolescent development. Some studies have shown that PFC neuroanatomical development correlates with levels of pubertal gonadal hormones (Herting et al., 2015; Nguyen et al., 2012; Peper et al., 2009; Perrin et al., 2009), though a complex picture of the interactive effects of pubertal hormones, cortical development and executive functioning has emerged in the literature (Hidalgo-Lopez & Pletzer, 2017; Stoica et al., 2019). Nonetheless, the continual development of the adolescent PFC combined with increasing cognitive ability suggests that the adolescent frontal cortex is in a state of heightened plasticity.

Male and female rats undergo similar cellular remodeling within the rodent medial prefrontal cortex (mPFC), often coinciding with or dependent upon pubertal onset, especially in females (Drzewiecki et al., 2016; Koss et al., 2015; Willing & Juraska, 2015). Additionally, mPFC-dependent cognitive behaviors mature during adolescence, in particular those that involve behavioral inhibition and cognitive flexibility (Andrzejewski et al., 2011; Koss et al., 2011; Newman & Mcgaughy, 2011; Willing et al., 2016). Pubertal onset appears to contribute to cognitive maturation, as male and female rats that are just 2 days post-pubertal perform better on a cognitive flexibility task than their prepubertal counterparts (Willing et al., 2016), though more information on the influences of puberty on adolescent development is needed.

Increasing inhibition in the mPFC during adolescence could contribute to a sensitive period for cortical plasticity. While the density of PV positive cells in the mPFC reaches adult-like levels in the juvenile period (Baker et al., 2017; Brenhouse & Andersen, 2011; Mix et al., 2015), PV protein expression continues to increase across mid to late adolescence (Caballero et al., 2014). Given the role of PV in maintaining sensitive periods in other sensory cortices, this could imply that the adolescent mPFC is in a similar sensitive period that closes sometime in young adulthood when PNNs reach adult-like levels between P60 and 70 (Baker et al., 2017).

However, further examination of the role of gonadal hormones on PNNs in males and females is warranted given that pubertal onset often coincides with or directly influences adolescent frontal cortex anatomical and functional development (e.g. Drzewiecki et al., 2016; Koss et al., 2011; Piekarski et al., 2017; Willing et al., 2016). Additionally, because the adolescent mPFC has been shown to undergo volumetric decreases between the juvenile period and adulthood (Markham et al., 2007), PNN density could be confounded by changes in volume, and a stereological approach for PNN quantification is necessary. Thus, this study was designed to stereologically quantify the total number of PNNs in the mPFC as well as frontal white matter volume as a broad indication of myelination across various stages of adolescence while accounting for pubertal status in both male and female rats.

## Methods

### Subjects

Subjects were the offspring of Long Evans rats ordered from Envigo [Formerly Harlan] (Indianapolis, IN) that were bred in the vivarium of the Psychology Department at the University of Illinois. Rats were weaned at P25, housed in groups of two or three with same-sex littermates and provided food and water *ad libitum* with a 12:12 light-dark cycle. All procedures were approved and adhered to guidelines set forth by the University of Illinois Institutional Care and Use Committee.

Pubertal status was assessed daily beginning at P29 in females and P39 in males due to the sex differences in pubertal timing. Among females, pubertal onset was determined by vaginal opening, which corresponds with increases in luteinizing hormone and estrogen (Castellano et al., 2011). In males, pubertal onset was identified by the separation of the prepuce from the glans penis, a process dependent upon the increased level of androgens at puberty (Korenbrot et al., 1977).

Rats were sacrificed at one of four timepoints. Both males and females were sacrificed at P30 and P60, ages that represent early adolescence and early adulthood, respectively. The other timepoints for sacrifice in this study were determined based on pubertal status. Within a litter, when a subject reached puberty, it was sacrificed 24 hours later with an age- and sex- matched sibling that had not yet reached puberty. An age-matched sibling of the opposite sex was also collected at this time point. These groups are referred to as the P35 and P43 groups, though it should be noted that these are the average ages of animals within each group.

All subjects were sacrificed between 10:00 and 15:00 to account for influences of circadian rhythms on PNN expression. For perfusion, subjects were deeply anesthetized with a lethal injection of sodium pentobarbitol and perfused intracardially with a 0.1 M solution of phosphate buffered saline (PBS) followed by a 4% paraformaldehyde fixative solution in PBS (PH 7.4). Brains were post-fixed in the paraformaldehyde solution for an additional 24 hours followed by immersion in a cryoprotectant 30% sucrose solution. At this point all brains were coded to conceal group from the experimenters. After 72 hours, brains were sliced in 40 µm coronal sections using a freezing microtome. Sections were stored at −20°C in an anti-freeze storage solution (30% glycerol, 30% ethylene glycol, 30% dH_2_0, 10% 0.2 M PBS).

### Staining with Methylene Blue/Azure II

Every other section containing the mPFC was washed with a 0.1 M solution of PBS and mounted on electrostatically charged slides. Once dry, slices were stained with the cell body stain Methylene Blue/Azure II and coverslipped with Permount.

### Volume quantification of the mPFC and white matter

The volume of the mPFC was quantified using the sections stained with Methylene Blue/Azure II and Stereoinvestigator software (Microbrightfield). The methodology has been previously described by our laboratory (Markham et al., 2007; Willing & Juraska, 2015). The dorsal and ventral boundaries of the mPFC were demarcated by an investigator blind to the age and sex of each subject. Briefly, the dorsal border of the mPFC was marked by decreased neuron density and altered laminar patterns in layers V/VI, and a less-defined cellular border between layers I and II. The ventral border of the mPFC was determined by decreases in cellular lamination of all layers. The mPFC was also divided into two subregions which contained the prelimbic cortex (PL, alternatively Cg3) and the infralimbic cortex (IL); the distinction between these two regions was determined by the lamination between layers III and V. Layer I was not included in the final analysis as it did not contain any neurons with visible PNNs (Fig.1).

**Figure 1.**
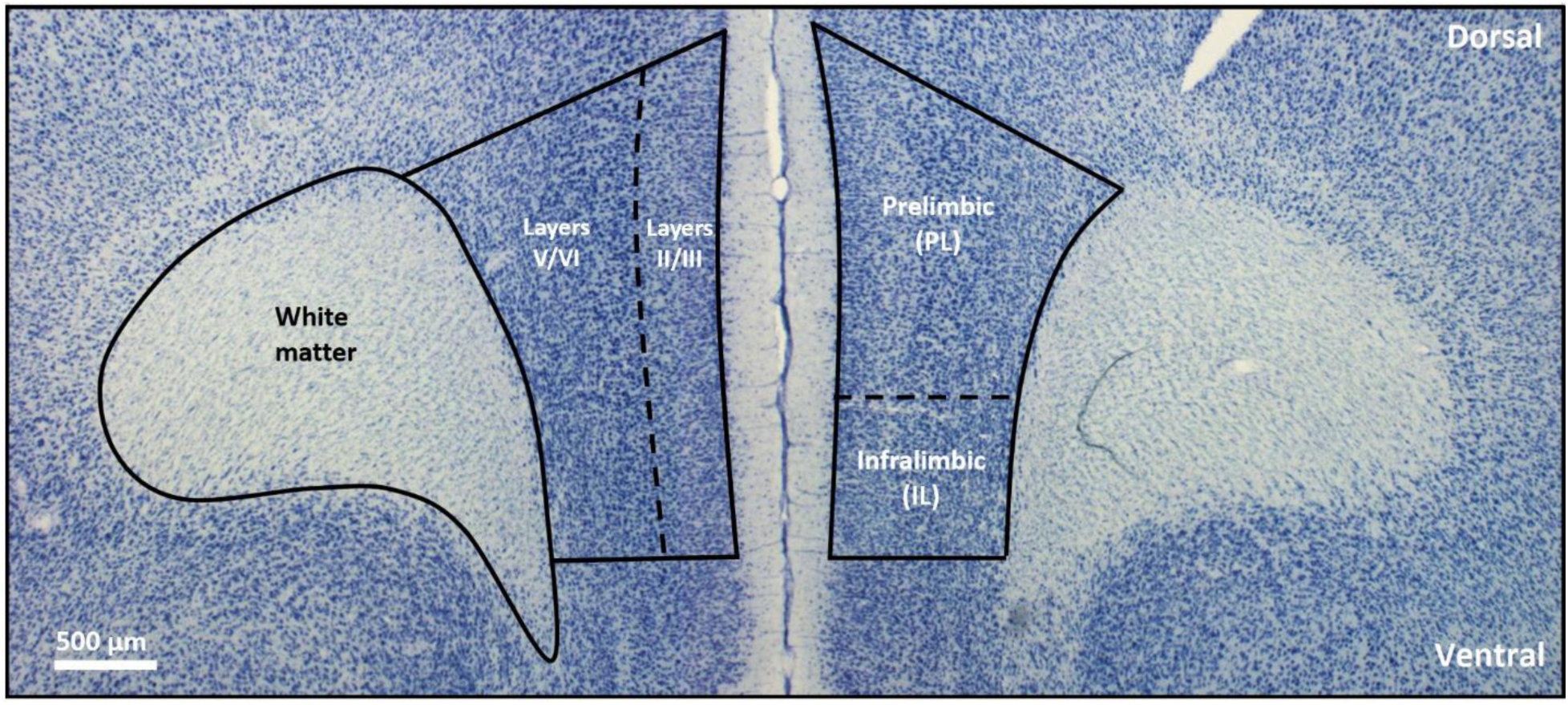
A coronal section of the mPFC stained with Methylene Blue/Azure II illustrating the delineation of white matter, layers II/II and V/VI, and the IL and PL subregions

Parcellation of the mPFC and white matter under it began on slices where the frontal white matter was first definitively present and ended on the caudal slices where the genu of the corpus callosum appeared. On average, 14 slices were parcellated for each subject. The post-fixative tissue thickness was also measured across the z-axis by determining the depth of the focal plane between the top and the bottom of clearly visible tissue. At least 50 thickness measurements were collected for each subject. The average thickness was multiplied by the total parcellated area to calculate to total volume of the gray and white matter in the mPFC. *Visualization of* Wisteria floribunda *agglutinin.* Every eighth section containing mPFC was stained with the biotinylated lectin *Wisteria floribunda* agglutinin (WFA, Sigma Aldrich, catalog L1516). This lectin reacts with the CSPGs and is commonly used as one of the broadest markers for PNNs (Giamanco et al., 2010; Härtig et al., 1994). Staining began by rinsing the tissue in triplicate with a tris buffered saline solution (TBS), followed by a 30 minute incubation at room temperature in a blocking solution (1% H_2_0_2_, 20% normal goal serum [NGS], 1% bovine serum albumin [BSA] in TBS) to prevent endogenous peroxide activity. Following blocking, the tissue was incubated for 24 hours at 4°C in WFA (1:500 concentration), which was diluted in Tris-Triton goat (TTG) solution (2% NSG, 0.3% Triton X-100 in TBS). The following day, sections were rinsed twice with both TTG and TBS, followed by a 60 minute incubation in avidin-biotin complex (ABC, Vectastain ABC Kit, Vector Laboratories) and triplicate 5-minute TBS rinses. Finally, slices were placed into a diaminobenzidine (DAB) solution in dH_2_0 (Sigma-Aldrich Fast 3–3’ Diaminobenzidine Tablets) for 2 minutes and then rinsed thoroughly with TBS to prevent further DAB reactions. Brain slices were mounted on electrostatically charged slides and allowed to dry for 24 hours before coverslipping with Permount (Fig.2a).

**Figure 2.**
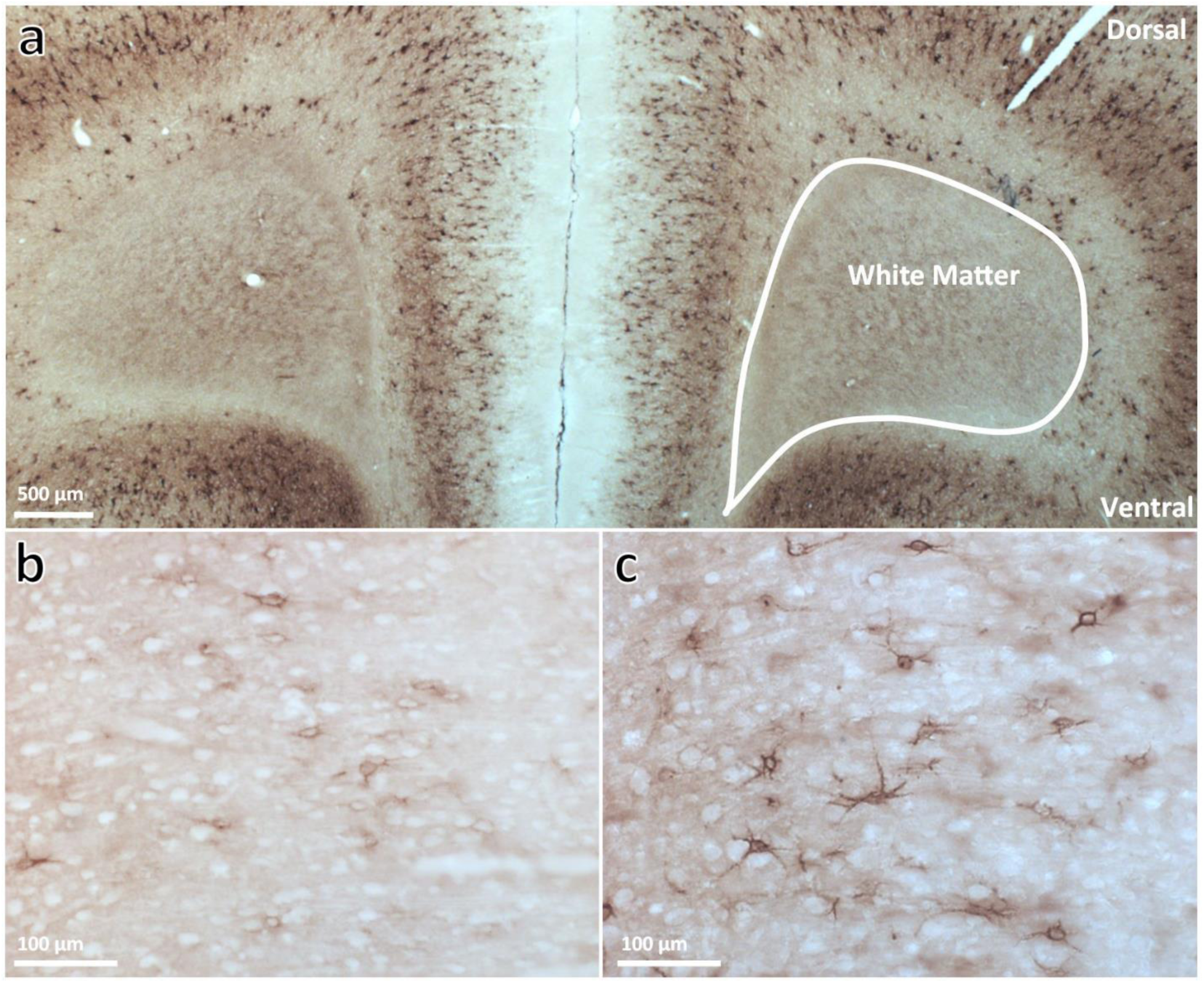
PNNs visualized with WFA in a coronal section of the mPFC. Shown across the left and right hemispheres (a) and at a higher magnification in layer V/VI of the mPFC (b, c). Both PNN staining and background intensity increased between P30 (b) and P60 (c).

### PNN quantification

The number of cells surrounded by a PNN were manually counted from two WFA stained sections using the StereoInvestigator optical disector described above. An experimenter blind to the age and sex of each animal manually traced a sampling area over the PL and IL regions. Layer I was not included in the sampling area due to the lack of cell bodies. The StereoInvestigator software then randomly determined sampling sites to count from using a 100 x 100 x 20 µm (length x width x height) counting frame with 1 µm guard zones. The counting frame contained two “exclusion” and two “inclusion” lines, and cells with a PNN were counted if they were included in the volume of the counting frame. The average densities of PNNs were calculated separately for the PL and IL and then multiplied by the previously measured volumes (with the exclusion of layer I) to obtain the total number of PNNs. Given difficulties in parcellating the WFA stained tissue, layer-specific PNN density quantification was not possible (Fig.2b).

### Statistical analysis

All statistical tests were conducted using RStudio. Using the “lmerTest” package, each sex was analyzed separately because of the known differences in pubertal timing and the differing degree of pubertal effects between the sexes found previously (Drzewiecki et al., 2016; Willing & Juraska, 2015). Levene’s test for equality of variances using the “car” package was conducted for each variable analyzed to confirm homogeneity of variance between groups. A linear mixed effects model for PNN number and white matter volume was then run with litter included as a random factor. Separate tests for analysis of variance (ANOVA) were conducted on the models, and post hoc analysis was conducted using the “emmeans” package on all significant ANOVA models. Post hoc tests ran pairwise comparisons using a Tukey adjustment. Additionally, a pre-planned paired t-test was conducted on the age- and litter-matched pre- and post-pubertal siblings (P35 in females and P43 in males) using a Bonferroni correction for two multiple, pre-planned comparisons. A total of 45 males and 44 females were analyzed (n=8-9/group).

## Results

### Pubertal onset

For all subjects sacrificed after puberty, the average age of pubertal onset was 34.9 days in females and 42.33 days in males. This is consistent with previous findings from our laboratory (Kougias et al., 2018). Among the pre- and post-pubertal siblings, the average age at sacrifice was 34.55 days with a range of 32-37 for females and 43 days with a range of 41-45 in males.

### PNN Density

The density of PNNs across all ages in males (Table 1) and females (Table 2) are presented as the average number of cells with a PNN per 200,000 µm^3^ (the volume of the counting frame used).

**Table 1.** Male PNN Density. PNN density in male subjects across the mPFC and its subareas, presented as mean number of PNNs per counting frame volume ± SEM. [* = p < 0.05 when compared to P60]

|  | P30 | P35 | P43<br>Pre-pubertal | P43<br>Post-pubertal | P60 |
| --- | --- | --- | --- | --- | --- |
| mPFC | 0.543 ± 0.06 * | 0.599 ± 0.05 | 0.597 ± 0.05 | 0.643 ± 0.06 | 0.840 ± 0.08 |
| PL | 0.620 ± 0.06 | 0.653 ± 0.05 | 0.646 ± 0.06 | 0.722 ± 0.06 | 0.908 ± 0.10 |
| IL | 0.397 ± 0.06 * | 0.500 ± 0.06 | 0.510 ± 0.06 | 0.500 ± 0.06 | 0.715 ± 0.08 |

**Table 2.** Female PNN Density. PNN density in female subjects across the mPFC and its subareas, presented as mean number of PNNs per counting frame volume ± SEM. [* = p < 0.05 when compared to P60; ** = p < 0.01 when come pared to P60; *** = p < 0.001 when compared to P60; ┼ = p < 0.05 when compared to pre-pubertal siblings]

|  | P30 | P35<br>Pre-pubertal | P35<br>Post-pubertal | P43 | P60 |
| --- | --- | --- | --- | --- | --- |
| mPFC | 0.572 ± 0.07 ** | 0.733 ± 0.07 | 0.574 ± 0.03 *, † | 0.647 ± 0.06<br>* | 0.955 ± 0.11 |
| PL | 0.645 ± 0.08 * | 0.796 ± 0.06 | 0.634 ± 0.05 *, † | 0.708 ± 0.06 | 0.975 ± 0.11 |
| IL | 0.428 ± 0.06<br>*** | 0.605 ± 0.09<br>* | 0.459 ± 0.05 *** | 0.536 ± 0.05<br>** | 0.920 ± 0.12 |

### PNN Intensity

In both sexes the intensity of PNNs increased with age (Fig. 2 b,c), as has been observed by other investigators in males (Baker et al., 2017; Mix et al., 2015). We did not quantify PNN intensity because the background was very variable across cellular layers (Fig. 2a). Additionally, the background staining intensity increased with age which would make comparisons between ages invalid.

### PNN Number

*Males.* In males, there was a significant main effect of age on the number of PNNs across the mPFC (*F*_4,40_ = 3.43, *p* = 0.017). Post hoc tests indicated that there was a significant increase in the total number of mPFC PNNs between P30 and P60 (*p* = 0.014) and P35 and P60 (*p* = 0.043). A paired t-test between age-matched pre- and post-pubertal siblings found no effect of puberty on total PNN number in the mPFC (*p* = 0.55) (Fig.3a).

**Figure 3.**
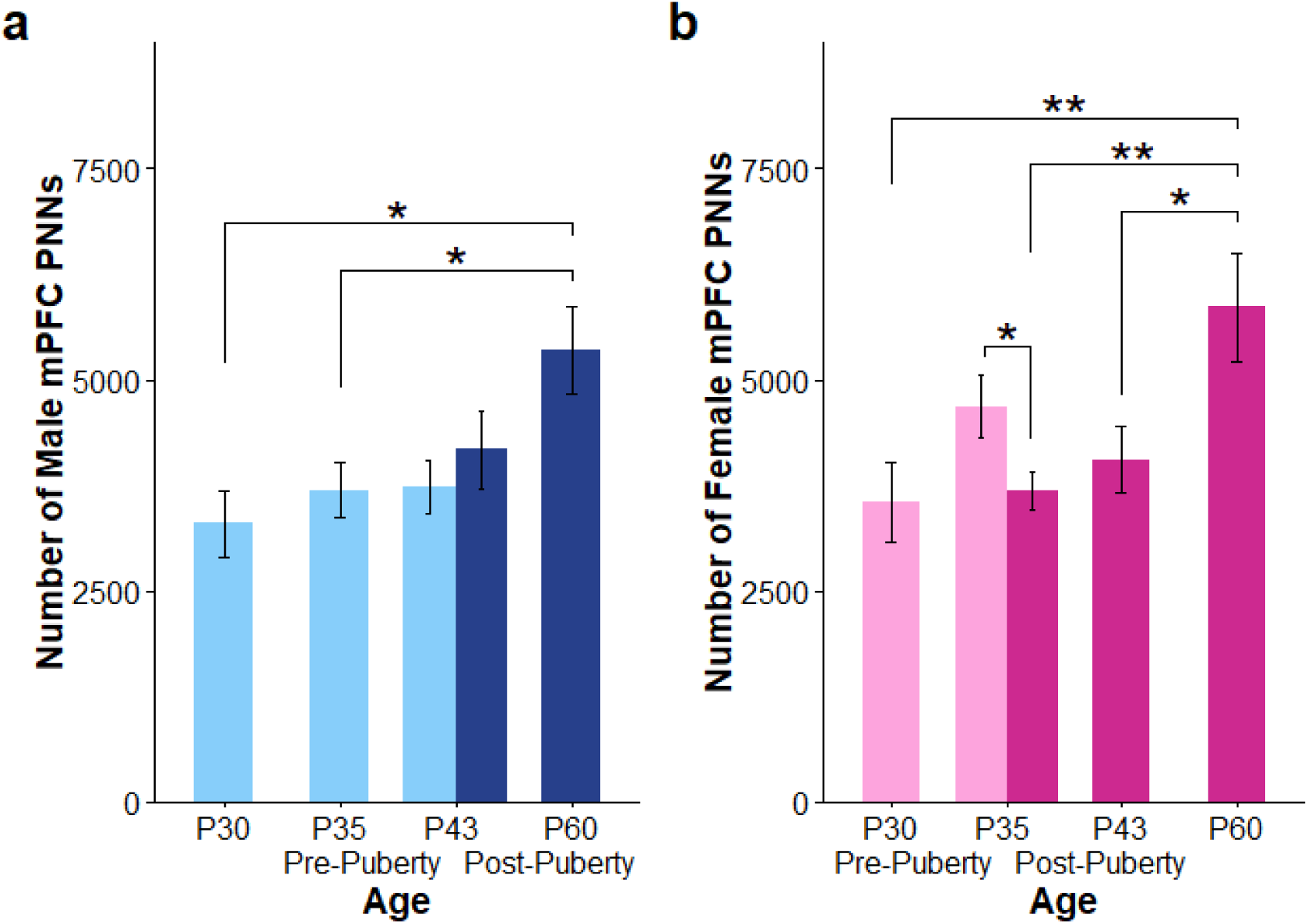
The number of mPFC PNNs in males (a) and females (b). There was a significant increase in the number of PNNs across the mPFC between P30 and P60 in both sexes. The lighter colored bars indicate animals that were pre-pubertal at sacrifice, while the darker bars indicate post-pubertal animals. In females, there was a decrease in the number of PNNs at puberty that persisted through P43 before reaching adult-like levels at P60. This effect of pubertal onset was not seen in males [* = *p* < 0.05; ** = *p* < 0.01]

There was a significant effect of age on the number of PL PNNs (*F*_4,40_ = 2.73, *p* = 0.042). Within this subarea, the number of PNNs increased significantly between P30 and P60 (*p* = 0.049) and showed a statistical trend when comparing P35 to P60 (*p* = 0.069). There were no effects of pubertal onset on PNN expression within the PL (*p* = 0.44) (Fig.4a).

**Figure 4.**
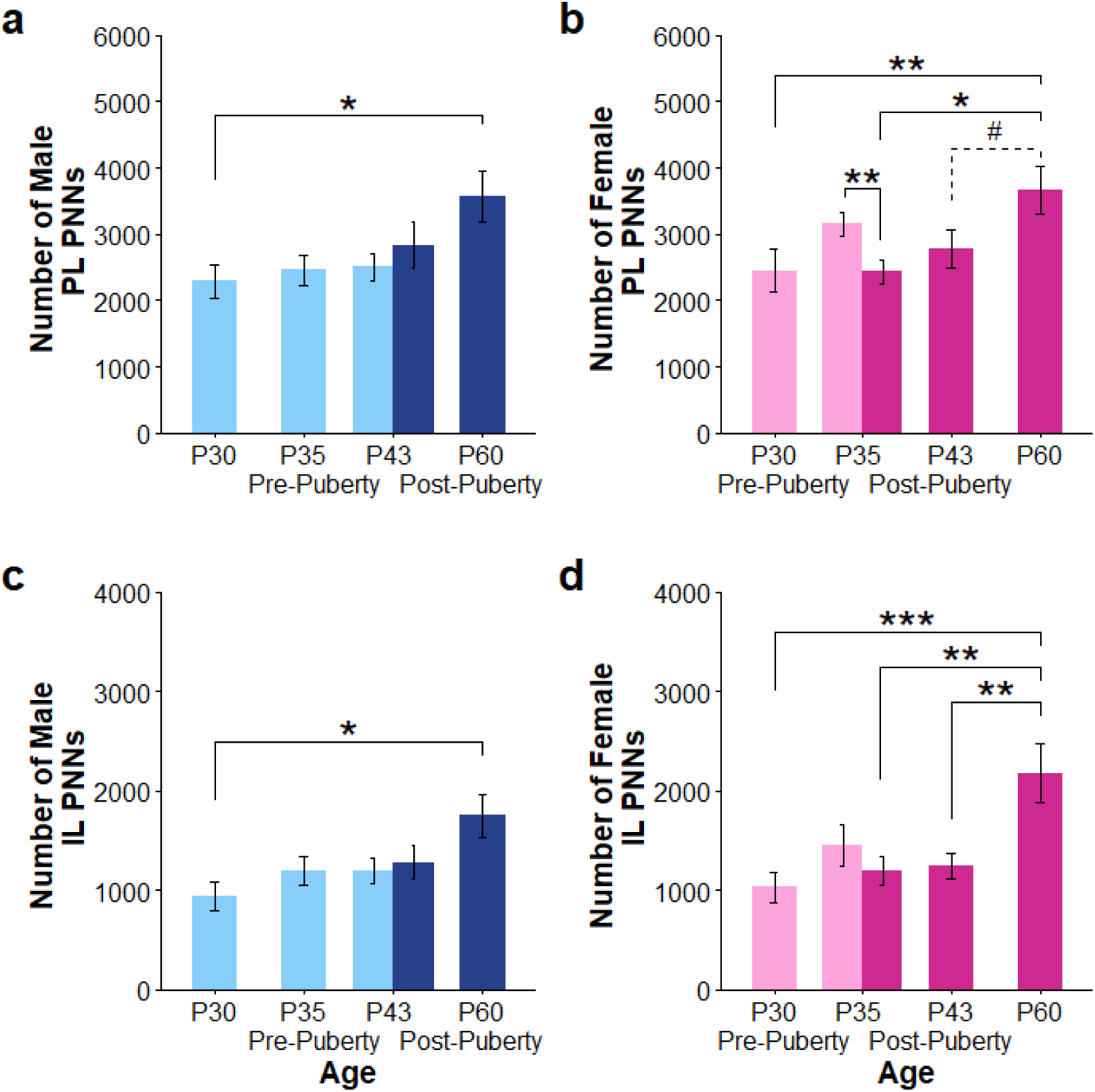
The number of PNNs in the PL and IL subregions. In males, PNNs in the PL (a) and IL (c) gradually increased between P30 and P60 with no influences of pubertal status on PNN number. A similar pattern was seen in the female PL (b) and IL (d), with a significant increase in PNNs occurring between P43 and P60. Pubertal onset was associated with decreased PNNs in the PL (b), as post-pubertal females had significantly fewer PNNs in this region compared to their pre-pubertal, age-matched siblings. This was not found in the IL (d) [# = *p* < 0.06; * = *p* < 0.05; ** = *p* < 0.01, *** = *p* < 0.001]

In the IL, there was also a main effect of age on PNN number (*F*_4,32_ = 2.94, *p* = 0.036). Post hoc analysis revealed a significant increase in IL PNNs between P30 and P60 (*p* = 0.018), with no influences of pubertal onset on PNN number (*p* = 0.95) (Fig.4c).

### Females

There was a main effect of age on the number of PNNs in the female mPFC (*F*_4,31_ = 6.31, *p* < 0.001). Post hoc analysis revealed that the number of PNNs across the mPFC increased significantly between P30 and P60 (*p* = 0.001). Additionally, the number of PNNs differed significantly between P35 post-pubertal females and P60 (*p* = 0.007) as well as the P43 and P60 females (*p* = 0.012). A paired t-test that compared pre-pubertal females to their age-matched, post-pubertal female siblings revealed a significant decrease in the total number of PNNs at pubertal onset (*p* = 0.02) (Fig.3b).

Within the PL subregion, there was a significant effect of age on PNN number (*F*_4,31_ = 5.04, *p* = 0.003). The number of PNNs in this region significantly increased between P30 and P60 (*p* = 0.008) and between post-pubertal P35 females and adults (*p* = 0.019). The difference between P43 and P60 females approached but did not reach statistical significance (*p* = 0.059). Pubertal onset significantly decreased PNNs in this region (*p* = 0.004) (Fig.4b).

There was also a main effect of age on the number of IL PNNs (*F*_4,32_ = 6.72, *p* < 0.001). Post hoc testing demonstrated that the total number of PNNs increased between P30 and P60 (*p* < 0.001), post-puberty P35 and P60 (*p* = 0.004), as well as P43 and P60 (*p* = 0.003). There was no effect of pubertal onset on PNNs in this region (*p* = 0.29) (Fig.4d).

### White Matter Volume

As expected, both males and females increased total white matter volume between P30 and P60. In males, there was a main effect of age on total white matter (*F*_4,911_ = 7.863, *p* < 0.001). Post hoc analysis demonstrated that white matter volume significantly increased between P30 and P60 (*p* < 0.0001) and between P35 and P60 (*p* < 0.001). White matter also significantly increased when comparing P30 males to P43 post-pubertal males (*p* = 0.0167) and trended toward statistical significance when comparing P30 males to P43 pre-pubertal males (*p* = 0.061). There were no significant increases in white matter volume between either P43 pre- or post-pubertal group and P60 subjects (*p* = 0.130 & *p* = 0.157, respectively), and no effects of pubertal onset on white matter volume (*p* = 0.999) (Fig.5a).

**Figure 5.**
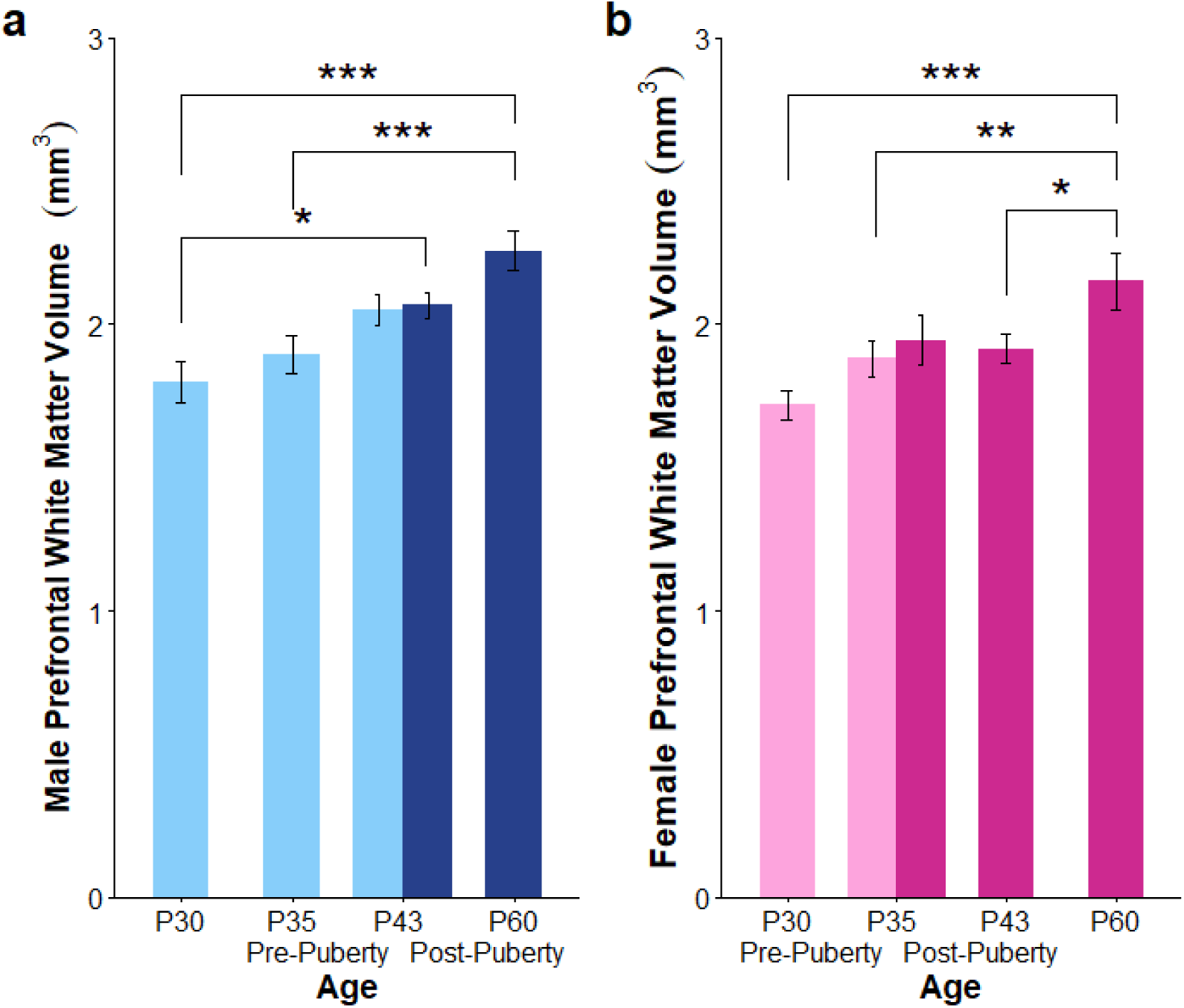
The volume of prefrontal white matter across adolescence in males (a) and females (b). Males reached young adult white matter levels by P43, while females showed a more protracted development, reaching adult-like white matter levels by P60. There were no significant effects of puberty on white matter volumes in males or females [* = *p* < 0.05; ** = *p* < 0.01, *** = *p* < 0.001]

Among females, there was a significant effect of age on white matter volume (*F*_4,533_ = 6.157, *p* < 0.001). The white matter significantly increased between P30 and P60 (*p* < 0.0001). There was significantly more white matter in the P60 females when compared to the pre-pubertal P35 females (*p* = 0.009) and the P42 females (*p* = 0.032), and the difference between P60 and P35 post-pubertal females trended toward statistical significance (*p* = 0.061). There was no significant effect of puberty on white matter volume (*p* = 0.951) (Fig.5b).

## Discussion

This study assessed both age-related changes and the influences of puberty on the number of PNNs in the developing adolescent mPFC in male and female rats. The number of PNNs increased between P30 and P60 in both sexes, similar to findings reported by Baker et al. (2017) in male rats. We also found a relationship between female pubertal onset and the number of PNNs, as pubertal onset was associated with an abrupt decrease in PNNs among females in the same litter that persisted for at least a week. This finding was somewhat unexpected, as previous anatomical work from our laboratory shows that post-pubertal females after P35 have an adult-like mPFC in terms of neuron and synapse number (Drzewiecki et al., 2016; Willing & Juraska, 2015). This protracted development of PNNs may result in prolonged cortical plasticity and contribute to the development of cognitive function during adolescence.

The sensitive period in the visual cortex opens with the appearance of the PV-expressing, inhibitory GABAergic cells and closes during the continual development of PNNs (Hensch, 2005; Pizzorusso et al., 2002). A similar anatomical pattern has emerged in the mPFC; PV expression becomes detectable during the juvenile stage and increases into adulthood (Caballero et al., 2014), and we have demonstrated an increase in PNNs in males and females during this time. As such, adolescence may be a sensitive period for the mPFC when the effects of experience are especially significant. For example, compared to juveniles and adults, adolescents are notably susceptible to the effects of social isolation (Einon & Morgan, 1977), which has long-lasting effects on mPFC function (Baarendse et al., 2013).

Altered adolescent plasticity in the mPFC could have implications for fear learning and extinction, which require prefrontal inhibitory/excitatory balance for top-down control of limbic structures (Santini et al., 2008; Sierra-Mercado et al., 2011). Degradation of mPFC PNNs in adult rodents can impair fear memory recall (Hylin et al., 2013), and we found low levels of PNNs during adolescence in both sexes. Adolescent male mice and rats do show attenuated fear extinction retention compared to juveniles and adults (McCallum et al., 2010; Pattwell et al., 2012). It is especially notable that male adolescent rats do not show deficits compared with adults on initial contextual fear learning and extinction, but unlike adults, they do show impaired memory for extinction 24 hours later that can be rescued with increased extinction training (Kim et al., 2011). Thus, both male mice and rats show fear memory deficits in adolescence when PNNs have not reached adult-like levels. However, it is unclear if the same impairments occur in adolescent females. Increased gonadal hormones in naturally-cycling adult females enhance extinction learning (Milad et al., 2009). Given this and the rapid decrease of mPFC PNNs we observed at pubertal onset, it is plausible that females would experience altered extinction memory at puberty.

One of the ways by which PNNs limit cortical plasticity is through physical and chemical barriers around neurons that prevent new synaptic contacts, as well as limiting AMPA receptor trafficking to the membrane (Frischknecht et al., 2009; Sorg et al., 2016). We have previously shown that female rats undergo synaptic pruning between P35 and P45 (Drzewiecki et al., 2016), coinciding with the decrease in PNNs observed here. It is possible that a downregulation of mPFC PNNs could be required for synaptic pruning to occur, and decreased PNNs during adolescence could also allow for dendritic remodeling in response to experience. For example, female rats with an induced lesion to the mPFC in early adolescence at P35 demonstrate greater dendritic remodeling than those who received the injury later in adolescence at P55 (Nemati & Kolb, 2012).

In addition to the development of PNNs, myelination can limit plasticity in the visual cortex through inhibitory interactions with the NgR, as demonstrated by McGee et al., (2005). Here, we measured the volume of the white matter under the prefrontal cortex. Unlike PNNs, there was not a difference in white matter volume in females that had reached puberty when compared to their pre-pubertal siblings. This is not surprising given the coarseness of simple volume measures. However, there was a pause in the increase in total white matter in females between P30 and P42 before rising again at P60. This is similar to what we reported in Willing et al. (2015), where females showed an interruption of growth at comparable ages while the white matter of males continued to increase linearly between P25 and P60. The underlying cellular basis may be the inhibitory influence of ovarian steroids on myelination, as pre-pubertal ovariectomy results in increases in the number of myelinated axons in the rat splenium assessed with electron microscopy (Yates & Juraska, 2008). The protracted pattern of white matter increases in the female mPFC could contribute to increased plasticity of this region during adolescence.

While the increase in PNNs appear to mark a decrease in experience-dependent plasticity, the functional significance of decreased PNNs at puberty in females is not entirely clear. PNNs have been linked to reversal learning capabilities, as degradation of PNNs or their components enhances reversal learning in adult male subjects (Happel et al., 2014; Morellini et al., 2010) and increased PNN expression in the mPFC is associated with decreased cognitive flexibility in adulthood (Coleman et al., 2014). Many studies have found that adolescent rats differ from adults on cognitive flexibility tasks (Koss et al., 2011; Newman & McGaughy, 2011; Simon et al., 2013; Westbrook et al., 2018), and we have found an increase in cognitive flexibility in both male and female rats two days after puberty (Willing et al., 2016). This increase in cognitive flexibility coincides with decreases in PNNs in females but not males. In addition, Willing et al. (2016) found the post-pubertal gains in cognitive flexibility continued into young adulthood, though the PNNs increased in late adolescence in both sexes here. Thus, there is not an obvious relationship between number of neurons surrounded by PNNs and the development of cognitive flexibility. Additionally, the influence of PNNs on cognition seems to be highly region and task-specific (e.g. Gogolla et al., 2009; Morellini et al., 2010; Romberg et al., 2013). As many aspects of cognition continue to mature across adolescence, the role of PNNs in learning and memory could differ across ages.

The abrupt decrease in PNNs at puberty in females suggests that estrogen may influence the maturation of inhibitory function in the mPFC. In particular, the estrogen receptor β (ERβ) colocalizes with the GABAergic PV cells throughout the adult cortex (Blurton-Jones & Tuszynski, 2002) and is highly expressed in the female mPFC (Shughrue et al., 1990). Additionally, the rise of pubertal estrogens is necessary for an increase in inhibition onto the excitatory neurons within the anterior cingulate cortex (Piekarski et al., 2017). However, we observed a decrease in mPFC PNNs at puberty in females, which may suggest a temporary decrease in inhibitory output. It is possible that estrogens act differently on the PNNs across various stages of development and in different cortical areas, though more research is needed to confirm this idea. Additionally, the number of PNNs in the male mPFC does not begin to increase until after pubertal onset, so we cannot exclude testosterone as a contributor to PNN formation. Furthermore, testosterone can be converted into estrogen via aromatase and can also activate estrogen receptors (e.g. Kawata, 1995), so work on the role for male gonadal hormones and PNN development is needed.

We observed similar PNN developmental trajectories in the PL and IL subdivisions of the mPFC in both sexes. In females, however, the post-pubertal decrease in PNNs was only significant in the PL, though a similar pattern was observed in the IL. This discrepancy could simply be due to the lower density and total number of cells surrounded by PNNs in the IL, requiring more power to detect a significant finding in this region. PNN expression throughout the cortex is highly region specific (Seeger et al., 1994; Ueno et al., 2017), so it is possible that they serve unique functions in each of the mPFC subdivisions. For instance, PNNs in the PL, but not IL, are required for acquisition and reconsolidation of cocaine memories in adult males (Slaker et al., 2015). Additionally, IL activation is required for fear extinction, which is enhanced under high estrogen conditions (Maeng et al., 2017; Milad et al., 2009; Sierra-Mercado et al., 2011), so it is possible that PNNs in the PL and IL respond differently to gonadal hormones.

Though we have previously shown that female rats lose approximately 12% of neurons in the mPFC around the time of pubertal onset, we do not attribute the observed decrease in PNN number here to this neuronal pruning. PNNs in this region primarily surround PV cells, and stereological analysis of GABA-ir positive cells in the mPFC revealed no differences between P35 and P45 females. In fact, this study notes a non-significant trend for an increase in the ratio of GABA-ir cells to the total mPFC neurons between P35 and P45, suggesting that proportionately more of the neurons pruned in adolescent females are excitatory (Willing & Juraska, 2015). Given that only 20-30% of PNNs surround non-PV cells in the mPFC (Baker et al., 2017; Slaker et al., 2015), it seems unlikely that this small subset of PNNs would be solely responsible for the 20% drop in PNNs we observed when comparing pre- and post-pubertal siblings.

Adolescence has been characterized as a period of vulnerability marked by emotional turmoil and stress (Casey et al., 2010; Crone & Dahl, 2012; Spear, 2000). The protracted development of mPFC PNNs observed here may contribute to this state of flux and potentially influence adolescent vulnerabilities to various psychiatric and substance use disorders (Chen et al., 2008; Kessler et al., 2005). However, it also could create a window of opportunity where stimulation and enrichment might be most beneficial. For example, PNNs have been shown to decrease in response to environmental enrichment, leading to an increase in plasticity (Sale et al., 2007). Because PNNs may be involved in aspects of psychiatric illnesses and drug seeking behaviors (Blacktop & Sorg, 2019; Mauney et al., 2013; Slaker et al., 2015), understanding PNN ontogeny could contribute to a better understanding of the mechanisms of adolescent vulnerability.

